# Reliable confidence intervals for RelTime estimates of evolutionary divergence times

**DOI:** 10.1101/677286

**Authors:** Qiqing Tao, Koichiro Tamura, Beatriz Mello, Sudhir Kumar

## Abstract

Confidence intervals (CIs) depict the statistical uncertainty surrounding evolutionary divergence time estimates. They capture variance contributed by the finite number of sequences and sites used in the alignment, deviations of evolutionary rates from a strict molecular clock in a phylogeny, and uncertainty associated with clock calibrations. Reliable tests of biological hypotheses demand reliable CIs. However, current non-Bayesian methods may produce unreliable CIs because they do not incorporate rate variation among lineages and interactions among clock calibrations properly. Here, we present a new analytical method to calculate CIs of divergence times estimated using the RelTime method, along with an approach to utilize multiple calibration uncertainty densities in these analyses. Empirical data analyses showed that the new methods produce CIs that overlap with Bayesian highest posterior density (HPD) intervals. In the analysis of computer-simulated data, we found that RelTime CIs show excellent average coverage probabilities, i.e., the true time is contained within the CIs with a 95% probability. These developments will encourage broader use of computationally-efficient RelTime approach in molecular dating analyses and biological hypothesis testing.

## Introduction

Reliable inference of the confidence intervals around the estimates of divergence times is essential for testing biological hypotheses (Burbrink and Pyron 2008; Kumar and Hedges 2016). Multiple sources contribute to the uncertainty of molecular divergence time estimates (Rannala and Yang 2007; Zhu et al. 2015; Kumar and Hedges 2016). One of them is the error associated with branch length estimation in a phylogeny due to the limited number of sites and substitutions in the sequence alignment (Kumar and Hedges 2016; Warnock et al. 2017). The stochastic nature of the substitution process (e.g., Poisson process) and the uncertainty involved in accounting for the unobserved substitutions (multiple-hit correction) result in errors in branch length estimates, which lead to imprecise time estimates (Kumar and Hedges 2016). However, this error decreases with an increase in the number of sampled sites (Rannala and Yang 2007; dos Reis and Yang 2013; Zhu et al. 2015) and becomes negligible for large phylogenomic datasets.

The second source of error is the variation of evolutionary rates among branches and lineages (Zhu et al. 2015; Kumar and Hedges 2016). Because rates and times are confounded, the variation of rates will naturally result in uncertainty of time estimates (Ho 2014; Zhu et al. 2015). This confounding effect cannot be eliminated by sampling more sites or genes in a dataset (Zhu et al. 2015; Kumar and Hedges 2016), so it contributes more uncertainty to time estimates than errors in branch length estimation for a large dataset. The uncertainty associated with clock calibrations due to the equivocal nature of fossil record presents a third source of error in divergence time estimation (Zhu et al. 2015; dos Reis et al. 2016; Warnock et al. 2017). The exact placement of fossil record in a phylogeny and the correct assignment of calibration constraints, especially the maximum constraint, are often difficult to justify, resulting in high uncertainty in the estimation of divergence time (Bromham et al. 2018).

In Bayesian analyses, the highest posterior density (HPD) intervals usually represent the uncertainty of inferred divergence times (Drummond et al. 2006). Bayesian methods compute HPD intervals directly from the density distribution of posterior times estimated using priors for branch rate heterogeneity, substitution process and fossil calibrations (dos Reis et al. 2016; Bromham et al. 2018), so sources contributing to the uncertainties of time estimates are automatically incorporated in the HPD intervals. Currently, Bayesian HPD intervals are considered reliable estimates of uncertainties surrounding divergence time estimates, although they are not always the same as the 95% confidence intervals (CIs) in the frequentist statistics (Jaynes and Kempthorne 1976; MacKenzie et al. 2017). Unfortunately, the enormous computational burden imposed by Bayesian approaches has hindered their applications to analyze many phylogenomic datasets (Pyron 2014; Mello et al. 2017; Li et al. 2019).

In contrast, non-Bayesian methods can analyze large-scale datasets quickly and generate accurate time estimates (Smith and O’Meara 2012; Tamura et al. 2012; Tamura et al. 2018). Unfortunately, the broad utility of these methods is still reduced by lack of reliable calculation of the uncertainty surrounding divergence times, which are represented by CIs. Non-Bayesian approaches require the use of analytical formulations or bootstrap approaches to estimate CIs (Sanderson 2003; Xia and Yang 2011; Tamura et al. 2013). However, site-resampling bootstrap approaches do not capture the error caused by rate heterogeneity, leading to false precisions of time estimates. Recognizing the need for incorporating lineage rate variation into CIs, Tamura et al. (2013) formulated analytical equations for the RelTime method, a non-Bayesian approach that relaxes the molecular clock. However, this approach may overestimate the amount of variance and produce overly wide confidence intervals (see below), resulting in low power for statistical testing (Kumar and Hedges 2016).

Bayesian and non-Bayesian methods also use different strategies to account for the uncertainty of fossil record. Non-Bayesian methods are currently limited to the use of minimum boundaries only, maximum boundaries only, or minimum and maximum boundary pairs as calibration constraints (Sanderson 2003; Tamura et al. 2013), while Bayesian methods allow the usage of probability densities as calibrations and automatically accommodate interactions among them (Inoue et al. 2010; Ho and Duchêne 2014). While Mello et al. (2017) presented a simple procedure to derive minimum and maximum boundaries from the density distributions, this strategy does not consider interactions among calibrations and may lead to overestimates of the variance of divergence times (see below).

Here, we present an analytical approach to estimate CIs for divergence times estimated using the RelTime method. The new analytical approach accounts for the variance associated with the branch lengths estimation as well as the variance due to rate heterogeneity to estimate CIs. We also present a simple approach to derive minimum and maximum boundaries from multiple calibration densities such that the calibration interactions are accommodated. Both approaches have been implemented in the MEGA X software for use in graphical and command-line interfaces (Kumar et al. 2012; Kumar et al. 2018). The 95% CIs produced by RelTime in empirical analyses are compared with the 95% HPD intervals produced by Bayesian methods to examine the performance of the new approaches. The approaches presented here may be used, with modifications, to improve variance calculation of time estimates for other non-Bayesian methods, e.g., penalized likelihood methods (Sanderson 2002).

## New Methods

### An analytical method to estimate confidence intervals

Considering a tree with three ingroup sequences (**Fig. 1**), relative time (*t*) for each node and relative rate (*r*) for each lineage are functions of branch lengths (*b*) in RelTime, e.g., *r*_1_, *r*_2_, *r*_3_, *r*_4_, *t*_*4*,_ and *t*_*5*_ are given by the following equations when the geometric means are used (similar equations can be derived when the arithmetic mean is used) (Tamura et al. 2018):

**Figure 1.**
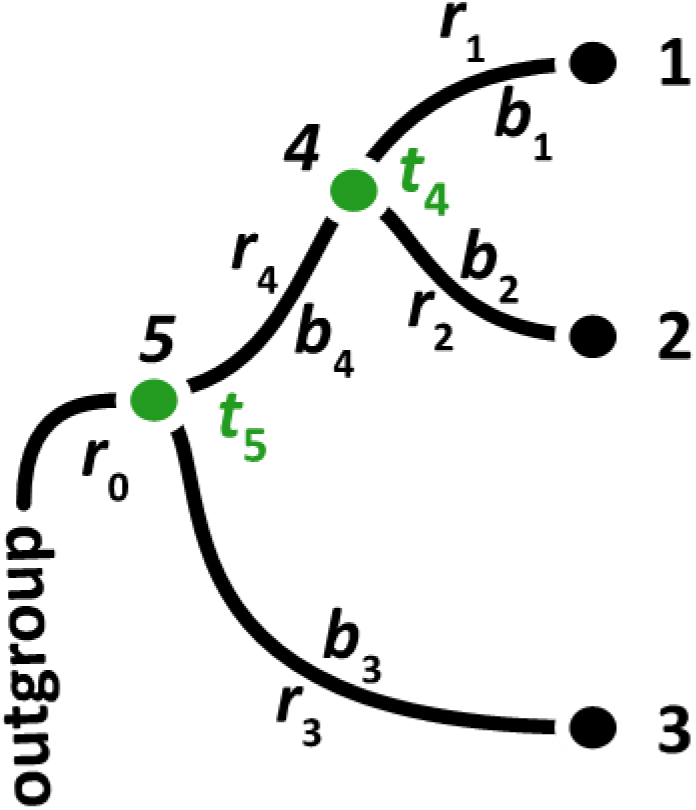
An evolutionary tree of three tips showing node times (*t*_i’_s), branch lengths (*b*_j_’s), and branch rates (*r*_j_’s).

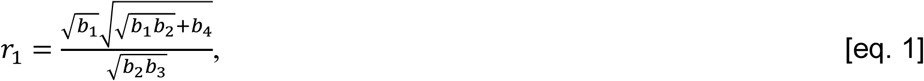

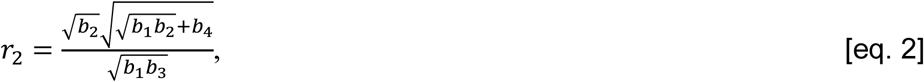

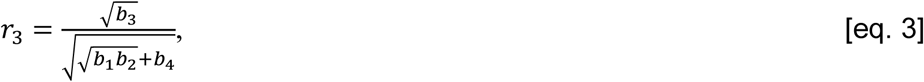

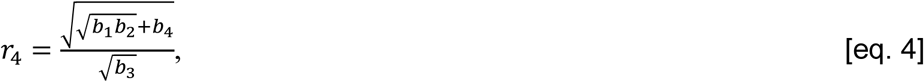

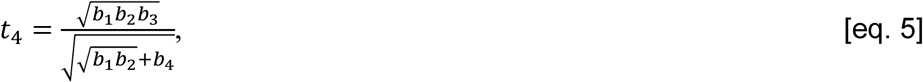

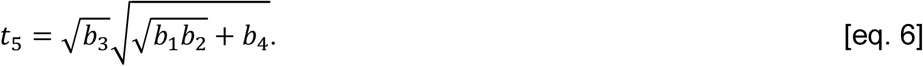

The variance of the estimated time (*t*_*i*_) for node *i*, denoted by *ν*(*t*_*i*_), can be estimated by the delta method, assuming that there is no covariance among branch lengths (*b*_*j*_’s):

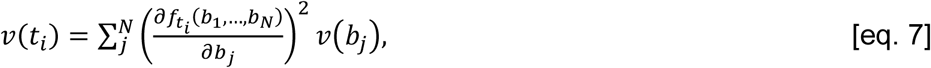

where *N* is the total number of branches, 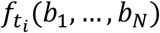 stands for the analytical function of *b*_*j*_’s to compute *t*_*i*_ (e.g., eq. 5 and eq. 6 for *t*_*4*_ and *t*_*5*_, respectively), and *ν*(*b*_*j*_) stands for the variance of branch length for branch *j*. Therefore, *ν*(*b*_*j*_) is required for computing *ν*(*t*_*i*_).

As mentioned before, the uncertainty of time is related to the number of sampling sites and the degree of rate heterogeneity. We consider the total variance of branch lengths, *ν*(*b*_*j*_), which is required to compute *ν*(*t*_*i*_), as a summation of the variance due to site sampling, *ν*_*s*_ (*b*_*j*_), and the variance due to rate heterogeneity, *ν*_*R*_ (*b* _*j*_):

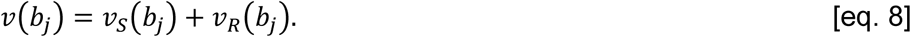

The value of *ν*_*s*_(*b*_*j*_) can be estimated by using analytical formulations or a site-resampling approach. For example, an approximate estimate of this variance can be obtained by the curvature method when the maximum likelihood method is used (Edwards 1992; Tamura et al. 2013).

However, it is more complex to estimate *ν*_*R*_ (*b*_*j*_), so we do it indirectly. We first compute the variance of observed evolutionary rates for all the lineages, *V*_*obs*_(*R*):

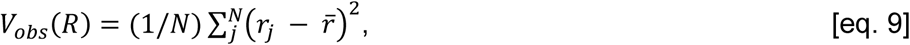

where R is a random variable representing all relative rates, *r*_j_ is the relative rate for each branch *j*, and 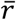 is the average of *r*_j_‘s. It is important to note that the relative rate for branch *j* is estimated as the relative rate for lineage *j* (Tamura et al. 2018). For example, RelTime computes the relative rate for *b*_4_ as the geometric mean of *r*_1_ and *r*_2_, which is assigned to be the rate for lineage *l*_4_ in **Figure 1**.

The variance of observed rates includes not only the variance introduced by rate heterogeneity, *R*_*V*_(*R*), but also the sampling variance associated with the branch length estimation, *S*_*V*_(*R*), because the observed relative rate *r*_j_ is calculated from branch lengths (*b*_*j*_’s) (e.g., equations 1 - 4). So,

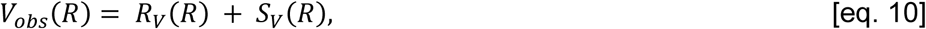

The value of *S*_*V*_(*R*) is obtained by summing the sampling variance of relative rate *r*_j_ for each branch *j*, denoted by *s*_*ν*_(*r*_*j*_):

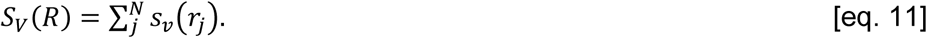

*s*_*ν*_(*r*_*j*_) can be estimated by the delta method, assuming that there is no covariance among *b*_*j*_’s:

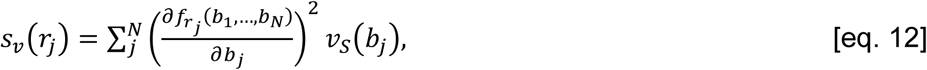

where 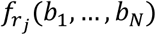 stands for the analytical function of *b*_*j*_’s to compute *r*_*j*_ (e.g., equation 1, 2, 3 and 4 for *r*_1_, *r*_2_, *r*_3_, and *r*_4_, respectively).

Using equations 9 – 12, we compute the variance introduced by rate heterogeneity:

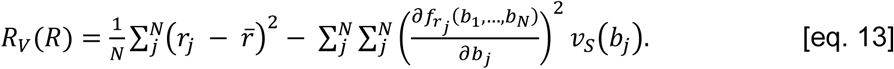

Then, we can compute the rate heterogeneity variance for each branch *j* as being proportional to its branch length:

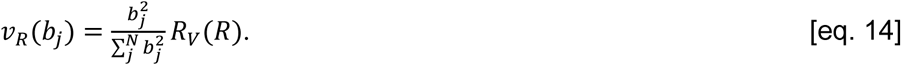

Using equations 8, 13 and 14, we can compute the total variance of branch length for branch *j*, denoted by *ν*(*b*_*j*_). Then *ν*(*b*_*j*_) can be used to compute the variance of time, *ν*(*t*_*i*_), using equation 7. For example, the variance of *t*_4_ and *t*_5_ are given by the following equations:

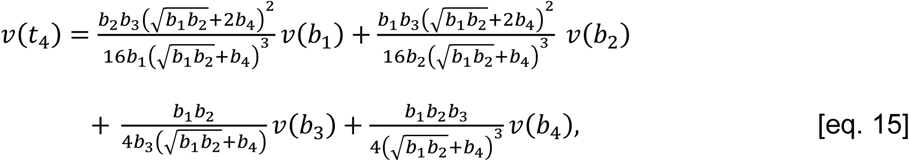

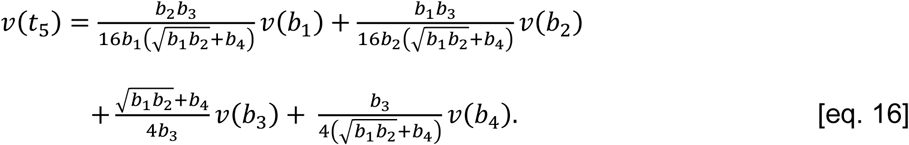

For larger numbers of taxa, such analytical formulations become complicated to derive, especially for deeper nodes. Thus, we compute the variance of divergence times for deeper nodes from tips to the root recursively. For example, using equations 15 and 16, we can derive

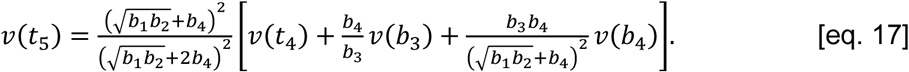

Therefore, the calculation of *ν*(*t*_5_) requires only *ν*(*t*_4_), *ν*(*b*_3_) and *ν*(*b*_4_), which are the variance for node *t*_4_ and branches *b*_3_ and *b*_4_, respectively. The variance of branches that do not directly connect to node 5, i.e., *ν*(*b*_1_) and *ν*(*b*_2_) in this case (**Fig. 1**) is not needed, if the value of *ν*(*t*_4_) is computed beforehand. Thus, for any node in a phylogeny, we can calculate the variance of divergence time recursively from tips to the root by using the variance of times for direct descent and ancestral nodes and the variance of directly connected branches. This procedure extraordinarily simplifies the computation of the variance of inferred time for each internal node in a tree with a large number of taxa.

It is important to note that times in the equations listed above are relative times, not absolute times because no calibrations are involved in the above equations. When one or multiple calibrations (minimum boundaries only, maximum boundaries only, or minimum and maximum boundary pairs) are given, RelTime will compute a global time factor (*f*) by altering relative times such that all calibration constraints are satisfied. When a range of *f* values can satisfy all calibration constraints, RelTime selects the midpoint of the range to be the best estimate of *f*. When one or more of the absolute times computed using the *f* value falls outside the calibration constraints, then RelTime adjusts relative times and *f* such that the deviations of absolute times from the calibration constraints are minimized. This process requires local alteration of relative rates and re-optimization of all other node times in the tree recursively (Tamura et al. 2013). For example, if the minimum age constraint of a node is violated, i.e., the age estimated using *f* is younger than the minimum constraint, RelTime decrease its estimate of the evolutionary rate proportionally in that lineage to adjust the age of this node to be higher, such that the divergence time becomes the same as the minimum age constraint. The resulting slowdown is transmitted to all the descendant nodes, and it affects the ancestral rates as well.

Similarly, if the maximum age constraint of a node is violated, i.e., the age estimated using *f* is older than the maximum constraint, RelTime increases the estimated evolutionary rate proportionally in that lineage such that the divergence time matches the maximum age constraint. Effects of this rate change will be transmitted to the descendant and ancestral nodes automatically. Consequently, RelTime will ensure that the absolute times for calibrated nodes are consistent with the user-desired calibration constraints.

In the final step, CIs are computed analytically using the final set of relative rates and the equations given above (e.g., equations 13-17), such that the uncertainty associated with clock calibrations can be incorporated into the CI calculation in RelTime. If the lower or upper bounds of CIs fall outside the user-specified calibration constraints, then CIs are truncated based on the imposed calibration constraints. Therefore, RelTime uses “hard” minimum and maximum bounds in CI calculation, as in BEAST (Bouckaert et al. 2014; Barba-Montoya et al. 2017).

### An approach to derive effective calibration boundaries from calibration densities

As stated above, calibration uncertainty is another critical source of estimation error in the inference of divergence times. Bayesian methods use various probability densities to accommodate the calibration uncertainty. However, the current non-Bayesian methods do not allow direct use of probability densities and do not provide provisions to incorporate interactions among calibration constraints. Therefore, we developed a new procedure for use in the RelTime method to derive calibration boundaries from probability densities that accounts for their interactions.

For each calibrated node with an associated probability density, we randomly sample two dates from the given probability density. We use these two sampled dates as the minimum and maximum (min-max) constraints for that node and derive such a min-max constraint for every node for which a probability density is specified. Then, we use all of these min-max boundaries to conduct RelTime analysis. We retain the RelTime time estimates only for the calibrated nodes, and then repeat the process of random sampling and dating for 10,000 times. A large number of iterations of this process ensure that calibration dates with tiny probabilities (0.01%) can be sampled.

The iterative procedure above produces a distribution of 10,000 inferred dates for each calibrated node. In the final step, we derive the minimum bound at 2.5% and the maximum bound at 97.5% of the distribution of inferred dates for each calibrated node. We refer to bounds derived during this process to be “effective bounds.” These effective bounds can be used together with the analytical approach described above to infer the divergence times and CIs in RelTime. It is important to note that effective bounds are used as calibration constraints, not densities. The actual shapes of the distribution of 10,000 inferred dates may vary slightly if one is to repeat 10,000 resamplings multiple times, but 2.5% and 97.5% boundaries of the distribution are expected to be stable, producing stable estimates of divergence times and CIs.

Our procedure is analogous to that in Bayesian methods, as both types of methods require resampling different sets of calibration constraints from user-specified densities, inference of divergence times using each set of sampled calibrations, and summarization of distributions of time estimates obtained from all sets of sampled calibrations. Therefore, the use of effective bounds allows RelTime to accommodate the interactions among calibration densities. However, it does not mean that RelTime and Bayesian methods are the same. Bayesian methods conduct calibration resampling and time inference steps simultaneously during the MCMC integration, whereas these steps are implemented sequentially in the RelTime method as proposed here.

We compared the effective bounds to calibration bounds derived using Mello et al. (2017)’s procedure (referenced as “Mello bounds” in the following) (**Fig. 2**), in which the minimum bound was placed at 2.5% of the density age, and the maximum bound was placed at 97.5% of the density age. Effective bounds were similar to the Mello bounds when the user-specified calibration density was reliable and informative, which meant that the true age of a node fell in the calibration density with a high probability. For example, effective bounds and Mello bounds almost overlapped for *Homo sapiens – Callithrix jacchus* split in which an exponential distribution was used as the calibration (**Fig. 2b**) (see the ***Materials and Methods*** section). When the user-specified density was uninformative, e.g., a diffused uniform distribution, Mello bounds were often diffused and matched the original density (**Fig. 2c**). In contrast, our new procedure generated narrower bounds due to the accommodation of the interactions among different calibration densities and constraints. These interactions reshaped the original, wider distribution and made it tighter (**Fig. 2c**).

**Figure 2.**
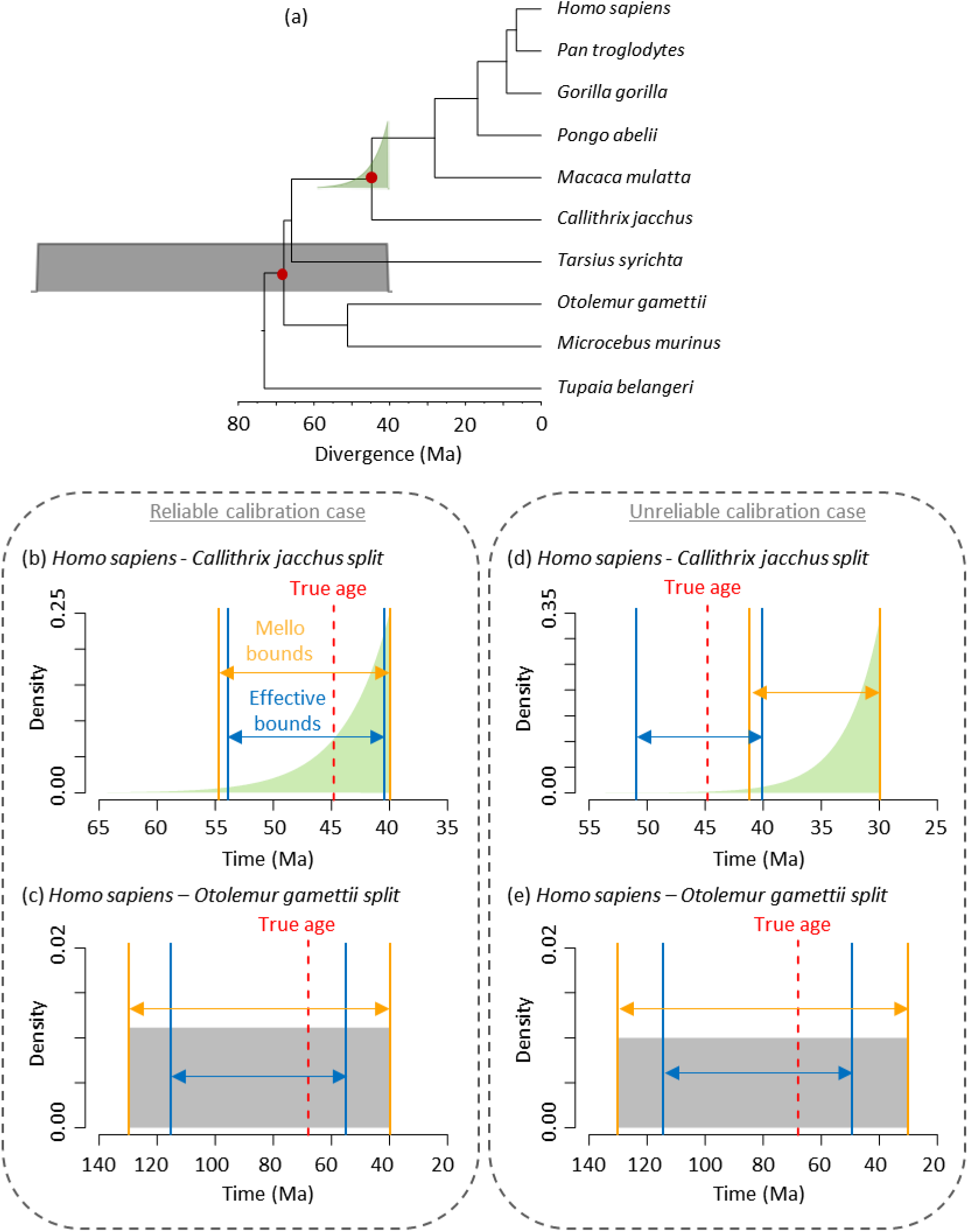
(a) A primate phylogeny with a user-specified uniform calibration density (gray shade) and an exponential calibration density (green shade). Red dots are the nodes shown in panels b-d. Effective bounds derived using our method (solid blue line) and bounds derived using Mello et al. (2017) procedure (solid orange line) are compared (b and c) when user-specified calibrations are reliable and (d and e) when user-specified calibration of *Homo sapiens – Callithrix jacchus* split is unreliable. The dashed red line represents the “true simulated age.”

Consequently, the use of effective bounds is likely to produce narrower CIs. In our analysis, when the user-specified calibration was unreliable, i.e., the true age of the node fell in its calibration density with a low probability, our effective bounds turned out to be better than Mello bounds. For example, when the true time of *Homo sapiens – Callithrix jacchus* split was located in the user-specified exponential density with < 2.5% probability (**Fig. 2d**), Mello bounds did not include the true time, resulting in incorrect time estimates. In contrast, our method did not ignore the low probability regions since it sampled 10,000 times from the user-specified density to ensure that dates with very low probabilities were considered. Thus, effective bounds are likely to contain the true time (**Fig. 2d**), and the use of effective bounds in RelTime may improve the accuracy and precision of time estimates.

## Results and Discussions

### RelTime produces CIs comparable to Bayesian HPD intervals in empirical analyses

We applied our methods to five empirical datasets containing nucleotide or protein sequence alignments from primates, spiders, insects, birds, and sun orchids (**Table 1**). We first present results from the primate dataset from Barba-Montoya et al. (2017), which contains a relatively small alignment of 9,361 base pairs from nine primate species and one outgroup (**Fig. 2a; Fig. S1a**). In this phylogeny, every internal node has been assigned a calibration density. Barba-Montoya et al. (2017) used two calibration strategies in MCMCTree (Yang 2007) and BEAST (Bouckaert et al. 2014) and compared the time estimates. We examined if the RelTime method produced estimates comparable to those obtained from Bayesian methods when all analyses employed the same alignment, phylogeny, substitution model, and calibration uncertainty densities (e.g., uniform distributions).

**Table 1.** Empirical datasets analyzed in this article.

| Taxonomic Group | Data type<sup>a</sup> | Sequence count<sup>b</sup> | Sequence length | Substitution model | Calibrations<sup>c</sup> | Software<sup>d</sup> | Reference |
| --- | --- | --- | --- | --- | --- | --- | --- |
| Primate (A) | N | 9 | 9,361 | HKY + G<sub>5</sub> | 8 uniform | MCMCTree<br>BEAST | Barba-Montoya et al. (2017) |
| Primate (B) | N | 9 | 9,361 | HKY + G<sub>5</sub> | 4 uniform<br>4 skewed | MCMCTree<br>BEAST | Barba-Montoya et al. (2017) |
| Spider | A | 40 | 55,447 | WAG + G<sub>5</sub> | 8 uniform | MCMCTree | Bond et al. (2014); Mello et al. (2017) |
| Insect | A | 143 | 220,091 | LG + G<sub>6</sub> | 38 uniform | MCMCTree | Tong et al. (2015) |
| Bird | N | 220 | 16,780 | HKY + G<sub>4</sub> | 13 uniform | BEAST | Oliveros et al. (2019) |
| Sun orchid | N | 57 | 773 | GTR + G<sub>4</sub> | 1 normal | BEAST | Nauheimer et al. (2018) |

<sup>a</sup>N = nucleotides; A = amino acids.
<sup>b</sup>Sequence count excludes the outgroup taxa.
<sup>c</sup>A Cauchy density distribution and an exponential density distribution are used as the skewed density in MCMCTree and BEAST, respectively.
<sup>d</sup>Software used in the original study for estimating divergence times.

For analyses of primate datasets where uniform densities were used as calibrations, we observed a high concordance between RelTime and Bayesian time estimates. The linear regression slopes were 0.97 and 1.03 when Bayesian analyses were conducted in MCMCTree and BEAST, respectively (**Fig. 3a** and **b**). This is a rather small difference. Although the width of RelTime CIs was slightly smaller than the width of Bayesian HPD intervals, RelTime CIs overlapped Bayesian HPD intervals for all the nodes (**Fig. 4a** and **b**). For primate datasets where a mixture of uniform and skewed densities were used as calibrations, RelTime estimates were again similar to Bayesian estimates, with a linear regression slope of 0.96 with MCMCTree (**Fig. 3c**) and 1.00 with BEAST estimates (**Fig. 3d**). RelTime CIs overlapped with MCMCTree and BEAST HPD intervals for all the nodes (**Fig. 4c** and **d**).

**Figure 3.**
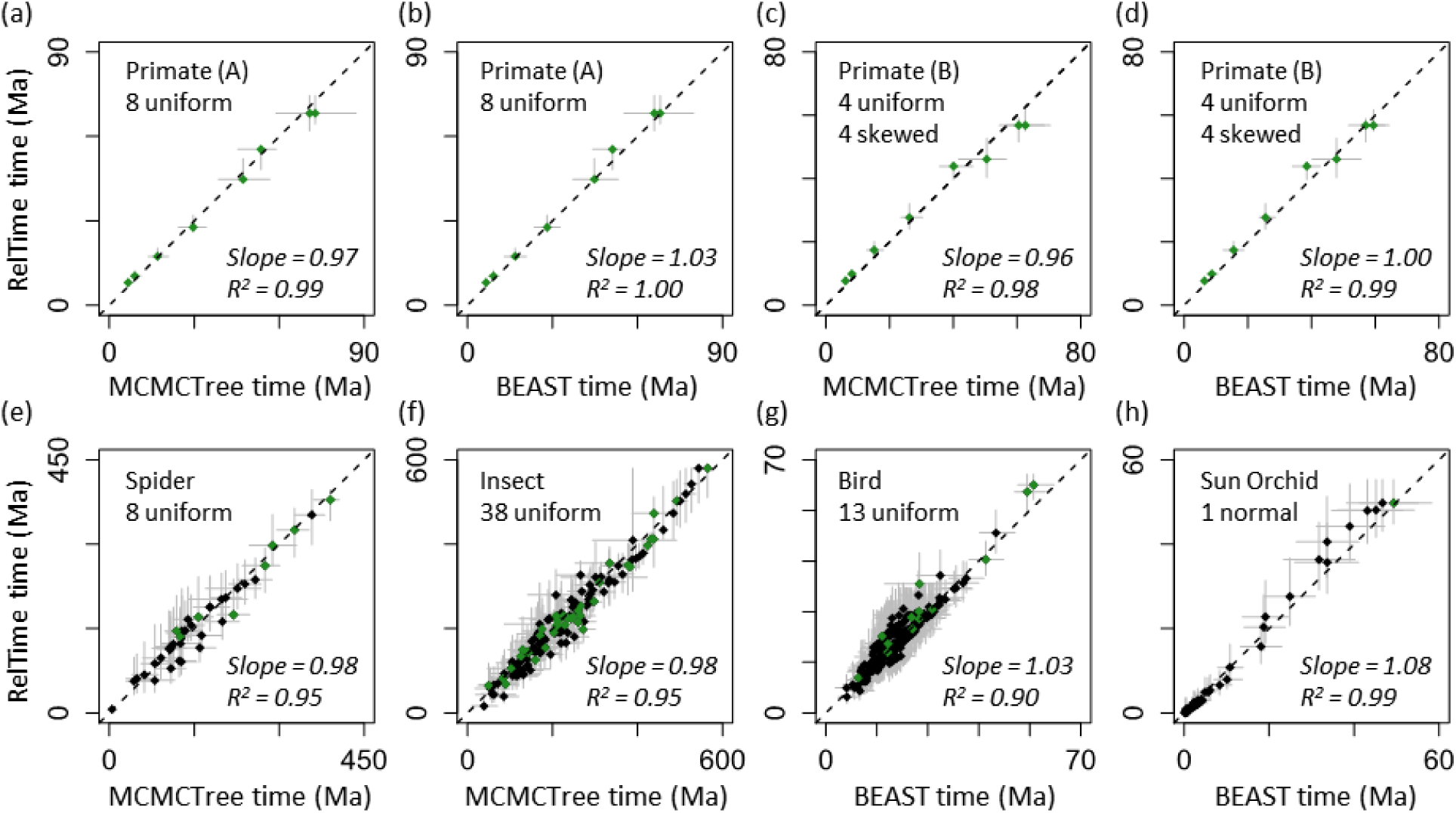
Comparisons of RelTime and Bayesian estimates of divergence times and the associated uncertainties. Gray bar represents the Bayesian 95% HPD intervals (x-axis) and RelTime 95% CIs (y-axis). Black dashed line represents 1:1 line. Each graph contains the slope and coefficient of determination (*R*2) values of the linear regression through the origin. Calibrated nodes are shown in green. The dataset name inside each panel refers to table 1.

**Figure 4.**
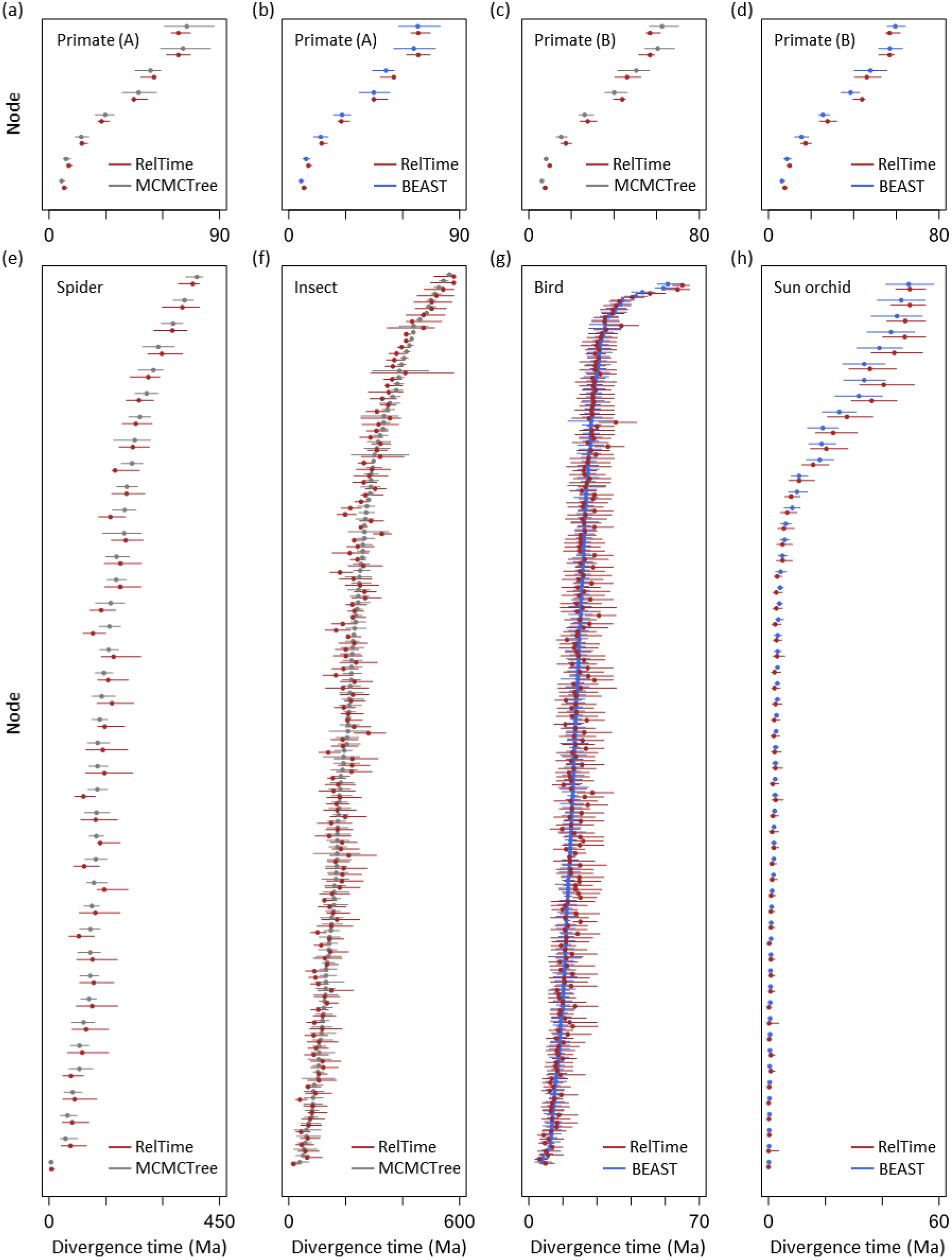
Comparisons of RelTime 95% CIs (dark red), MCMCTree 95% HPD intervals (gray), and BEAST 95% HPD intervals (blue). Dots are point estimates of divergence times. The dataset name inside each panel refers to table 1.

We then analyzed spider and insect datasets to examine the performance of our methods for larger datasets (>40 species and >50,000 sites). These datasets consisted of protein sequences and presented more extensive rate variation among branches and lineages as compared to the primate dataset (**Fig. S1b** and **c**). Fewer calibrations were used in these datasets with eight calibrated nodes in the spider dataset and 38 calibrated nodes in the insect dataset. This means that 20% - 26% nodes of the phylogeny were assigned calibration values. We again observed strong concordance between RelTime and Bayesian time estimates, with a linear slope of 0.98 and 0.98 for the spider and insect dataset, respectively (**Fig. 3e** and **f**). The high similarity between RelTime and Bayesian node times remained even after we excluded nodes on which user-specified calibrations were assigned (slope = 0.97 and 0.98, respectively). Although CIs produced by RelTime were slightly wider than HPD intervals produced by Bayesian methods, they were comparable with more than 97% of the nodes in spider and insect datasets showing overlapping CIs and HPD intervals (**Fig. 4e** and **f**). When CIs and HPD intervals did not overlap, they were less than 5 million years (Myr) apart. Therefore, RelTime CIs can be considered similar to Bayesian HPD intervals for these two datasets.

Although it is becoming more common to apply many internal calibrations to empirical studies, researchers may only have a limited number of calibrations because reliable fossil record for most taxonomic groups is limited. So, we analyzed another two nucleotide sequence alignments in which only a few calibrations have been used. One of them is a bird phylogeny (**Fig. S1d**) containing 220 ingroup taxa with only 13 calibrations (i.e., 6% nodes are calibrated). Again, RelTime produced time estimates similar to Bayesian estimates, showing a linear slope of 1.03 with slightly weaker linear relationship than seen for other datasets above (**Fig. 3g**). CIs produced by RelTime overlapped with HPD intervals provided by the Bayesian method for all the nodes except for two (**Fig. 4g**). For these two nodes, CIs and HPD intervals were less than 2 Myr apart. In the analysis of the sun orchid dataset, in which 57 sequences were included, and only a single internal calibration was used, time estimates obtained from RelTime and Bayesian methods showed a good linear relationship as well (**Fig. 3h**). Although RelTime generated slightly older time estimates than those from the Bayesian method, CIs and HPD intervals overlapped for all the nodes (**Fig. 4h**).

These results suggest that the application of our analytical method for computing CI combined with the approach to derive effective bounds is likely to produce estimates of times and CIs compatible with Bayesian estimates for datasets with a small and large number of calibrations when the same alignment, phylogeny, and calibration densities are used. We found that, compared to the previous implementations in RelTime for estimating CIs (Tamura et al. 2013; Mello et al. 2017), our new methods effectively improved the CI inference, because the width of CIs was reduced by 33% - 64% in the analysis of empirical data. This reduction was seen for both the calibrated and non-calibrated nodes. We attribute this improvement to the fact that the new analytical method accounts for the rate heterogeneity better and the effective bounds reflect calibration interactions and reshape the original diffused calibration densities to generate narrower CIs. Consequently, the precision of divergence time estimates is improved. However, it is essential to note that the use of incorrect calibration constraints or densities can significantly impact the precision of time estimates (Warnock et al. 2017). Therefore, one needs to examine the reliability of calibrations before conducting dating analyses (Andújar et al. 2014; Battistuzzi et al. 2015; Hedges et al. 2018).

### RelTime CIs show high coverage probabilities

We conducted RelTime and Bayesian (MCMCTree) analyses on a large set of simulation datasets, containing small and large numbers of sequences, to compare the overall coverage probabilities, i.e., the proportion of RelTime CIs and Bayesian HPD intervals that included the true divergence time. In these analyses, we used no ingroup calibrations to avoid confounding the effect of calibrations on the coverage probabilities of estimated CIs and HPD intervals (see the ***Methods and Materials*** section).

RelTime performed well in the analysis of datasets in which branch rates evolved under an independent branch rate (IBR) model, showing high coverage probabilities (≥ 95%) in both small and large simulated datasets (**Fig. 5a**). The Bayesian analyses also performed well for IBR datasets, showing high overall coverage probability for datasets containing 50 sequences (97%) but slightly lower overall coverage probability for datasets containing 100 sequences (84%) (**Fig. 5a**). Interestingly, the coverage probability of HPD intervals declined further for IBR datasets with 200 sequences (62%). We found that the true divergence times were located close to the boundaries of HPD intervals. In these cases, the average percent time difference between a true age and the nearest bound of respective HPD interval was 10%. RelTime showed an overall coverage probability of 94% for datasets containing 50 sequences and evolving with autocorrelated rates in a phylogeny (ABR), while the Bayesian method showed slightly lower coverage probabilities (78%; **Fig. 5b**). Bayesian coverage probabilities declined for datasets containing 100 and 200 sequences (52% and 36%, respectively), and average percent time difference between the true ages and their nearest HPD interval boundaries was large (20% - 45%). In contrast, RelTime maintained its high coverage probability for large datasets (≥ 95%).

**Figure 5.**
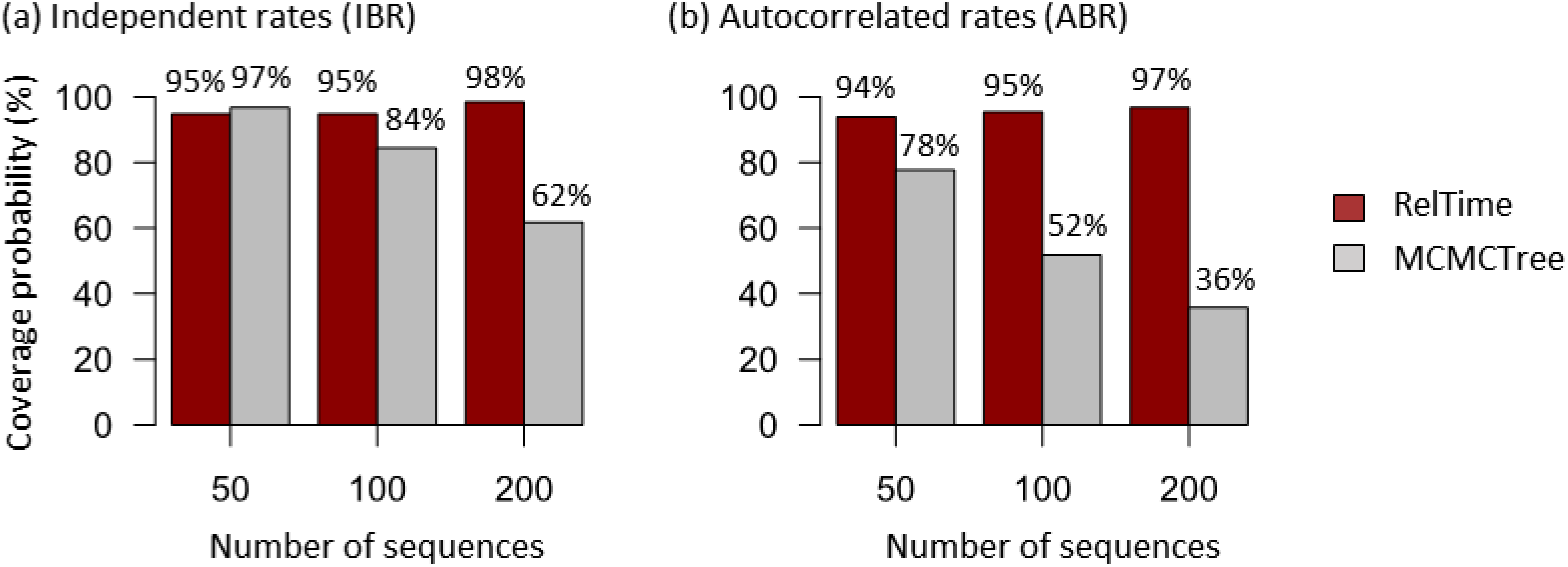
The overall coverage probabilities of RelTime CIs and Bayesian methods HPD intervals produced by analyzing datasets with different numbers of sequences simulated under an (a) independent branch rate, IBR, model and (b) autocorrelated branch rate, ABR, model.

We found that the posterior distributions of many nodes for ABR datasets were not normal-like and skewed, which means that the interpretation of HPD intervals is not really the same as the CIs, which has been noted earlier (Jaynes and Kempthorne 1976; MacKenzie et al. 2017). It is also known that the tree prior assumed in Bayesian analyses can have an impact on the estimation of divergence times and HPD intervals (Heled and Drummond 2014; Ritchie et al. 2017; Bromham et al. 2018). This impact is more prominent when there is limited number of calibrations because the tree prior provides node age priors for nodes without calibrations (Barba-Montoya et al. 2017), which may explain the observed low coverage probabilities. We expect that the Bayesian method will perform better if more informative calibrations are applied because the tree prior becomes less critical and informative calibrations reduce the uncertainty of time estimates. More extensive analysis of this problem is beyond the scope of this article, but we plan to pursue it in the future.

## Conclusions

Our new analytical method to estimate CIs, as well as the approach for deriving effective bounds, will now allow the use of more biological information, such as the rate variation among lineages and the probability density of calibrations, in 95% CIs inference. RelTime is computationally efficient, requiring only a fraction of the time and resources demanded by Bayesian approaches (Tamura et al. 2012; Tamura et al. 2018). Results from the analysis of empirical nucleotide and protein sequence alignments containing small and large numbers of sequences and calibrations suggest that RelTime will serve as a reliable approach for dating the tree of life and conducting biological hypothesis testing, especially for large-scale molecular data. We also anticipate that our new analytical method and new approach for utilizing calibration densities, with modifications, can be applied to generate CIs for other non-Bayesian dating methods, e.g., penalized likelihood methods (Sanderson 2002).

## Materials and Methods

### Comparisons of user-specified calibration density, Mello bounds, and effective bounds

We used the BEAST-generated primate timetree published in Barba-Montoya et al. (2017) as the true tree (**Fig. 2a**) and simulated an alignment of 9361 sites under HKY+G (Hasegawa et al. 1985) model in SeqGen with parameters derived from the empirical molecular data. Branch-specific rates were sampled from an uncorrelated lognormal distribution with a mean rate of 0.0069 substitutions per site per Ma and a standard deviation of 0.4 (log-scale). The simulated alignment was used to derive effective bounds.

We tested the performance of using effective bounds and Mello bounds under two calibration scenarios: reliable and unreliable scenarios. The reliable scenario represented the case where true ages of all the nodes are located in their calibration densities with high probabilities. The unreliable scenario refers to situations in which true ages of some nodes are found in calibration densities with low probabilities, whereas the true ages for some other nodes are found in calibration densities with high probabilities. An informative exponential density was used at *Homo sapiens – Callithrix jacchus* split (true age = 44.8Ma) and an uninformative uniform density were used at *Homo sapiens – Otolemur gamettii* split (true age = 68Ma) under both scenarios. In the reliable calibration scenario, we assumed that a minimum age of 40Ma at *Homo sapiens – Callithrix jacchus* split and maximum age of 130Ma at *Homo sapiens – Otolemur gamettii* split were known. Therefore, we used an exponential density (mean = 4Ma and offset = 40Ma) and a uniform density (min = 40Ma, max = 130Ma) at *Homo sapiens – Callithrix jacchus* split and at *Homo sapiens – Otolemur gamettii* split, respectively. The true ages of both nodes located in their densities with high probabilities. Under the unreliable calibration scenario, we assumed that a minimum age of 30Ma at *Homo sapiens – Callithrix jacchus* split and maximum age of 130Ma at *Homo sapiens – Otolemur gamettii* split were known. Therefore, we used an exponential density (mean = 3Ma and offset = 30Ma) and a uniform density (min = 30Ma, max = 130Ma) at *Homo sapiens – Callithrix jacchus* split and at *Homo sapiens – Otolemur gamettii* split, respectively. This resulted in the true age of *Homo sapiens – Callithrix jacchus* split located in its density with a low probability (< 2.5%), while the true age of *Homo sapiens – Otolemur gamettii* split located in its density with a high probability.

### Empirical analyses

We obtained six empirical datasets that employed different calibration strategies from five published studies (**Table 1**) (Bond et al. 2014; Tong et al. 2015; Barba-Montoya et al. 2017; Nauheimer et al. 2018; Oliveros et al. 2019). Molecular data were obtained from supplementary files of original studies. Calibration densities and Bayesian timetrees (including HPD intervals) were provided by authors or derived from the original studies, except for Bond et al. (2014)’s data, which was obtained from Mello et al. (2017). In RelTime analyses, we used the same alignments, substitution models, tree topologies, and calibration densities for ingroup nodes as the original studies to ensure comparability with Bayesian results. RelTime analyses were conducted in MEGA X (Kumar et al. 2018). For Oliveros et al. (2019)’s data, the published Bayesian timetree was summarized from 10 timetrees inferred using 10 different random subsets of the full dataset, because BEAST was computationally infeasible to analyze the full dataset. Since the original study has shown that 10 subsets provided similar results, we only conducted RelTime analysis using one subset. We compared RelTime time estimates and CIs with Bayesian time estimates and HPD intervals. We did not test whether the slope between RelTime and Bayesian time estimates was one because *p*-value will always reject the hypothesis of the slope of one when the data sample size is large, which makes its use less meaningful (Halsey 2019; Wasserstein et al. 2019). To compare the performance of our methods and the previous CI calculation methods for RelTime, we re-analyzed all empirical datasets using Mello bounds and the Tamura et al. (2013)’s method, which was implemented in MEGA 7 (Kumar et al. 2012; Kumar et al. 2016).

### Simulation analyses

We used the datasets simulated by Tamura et al. (2012), in which sequence alignments were generated using independent (IBR) and autocorrelated (ABR) branch rate models. In IBR cases, branch rates were sampled from a uniform distribution in the interval [0.5*r*, 1.5*r*], where r was the evolutionary rate derived from empirical genes. In ABR cases, branch rates were simulated using Kishino et al. (2001) model (lognormal distribution) with the initial rate of *r* and autocorrelation parameter *v* = 0.1 (time unit = 100Myr). GC contents, transition/transversion ratios, and sequence lengths were all derived from empirical genes and varied among datasets. Both ABR and IBR scenarios contained 100 simulated datasets, and each dataset contained 400 ingroup sequences. Because the Bayesian method requires a long runtime for analyzing a dataset with 400 sequences. We subsampled 50, 100, and 200 sequences from the original full datasets and conducted RelTime and Bayesian analyses for these subsets.

To examine the performance of RelTime on simulated datasets, we used the minimum number of calibrations, in order to avoid the possibility that the use of many informative calibrations mediated the similarity of performance of RelTime and Bayesian methods. In the Bayesian analysis, we used MCMCTree and a single calibration at the root with a diffused uniform density (true age ± 50 Myr). The use of diffused density could reduce the impact of calibration on constraining the width of HPD intervals. We used 100 Myr as the time unit and “rgene_gamma = 2 10” and “sigma2_gamma = 2 20” as priors, so the prior values of mean rate, independent rate variation, and autocorrelation parameter were similar to the true values. The use of lognormal distribution as the rate model in Bayesian analyses was appropriate because the lognormal distribution fit the distribution of evolutionary rates for IBR and ABR datasets, although IBR datasets were simulated using a uniform distribution. We used “BDparas = 2 2 0.1” as the tree prior because it generated a uniform node age prior and it was commonly used in practice (Yang 2006). Two independent runs of 100,000 generations each were conducted, and results were checked for good ESS values (>200) and convergence. In the RelTime analysis, we did not use any calibrations, so there was no calibration effect on constraining the width of CIs. However, because no calibration was used, RelTime provided relative times instead of absolution times. To make the fair comparison between RelTime and Bayesian results, we normalized the RelTime times (and CIs), Bayesian times (and HPD intervals) and true times to their ingroup root age, which was analogous to the case where the age of the ingroup crown node was fixed. We computed the coverage probability using these normalized times. The coverage probability of each node was the proportion of 100 datasets where the CI (or HPD interval) of this node contained the true time. Because the Bayesian method required a long computational time for analyzing a dataset with 200 sequences, we only finished the analyses of 50 datasets and used them to calculate the coverage probability for each node. The overall coverage probability was the mean value of the coverage probabilities of all the ingroup nodes.

## Acknowledgments

We thank Drs. Jose Barba-Montoya and Simon Ho for sharing Bayesian results, and Mary Kathleen Durnan and Joy Wenslas for the technical support. We also thank Drs. Sayaka Miura and Jose Barba-Montoya for critical comments. This research was supported by grants from the National Institutes of Health (NIH GM0126567-02), National Science Foundation (NSF 1661218), and National Aeronautics and Space Administration (NASA NNX16AJ30G) to SK, Brazilian Research Council (CNPq 233920/2014-5 and 409152/2018-8) to BM, and Tokyo Metropolitan University (DB105) to KT.

## Figure Legends

**Supplementary figure 1.**
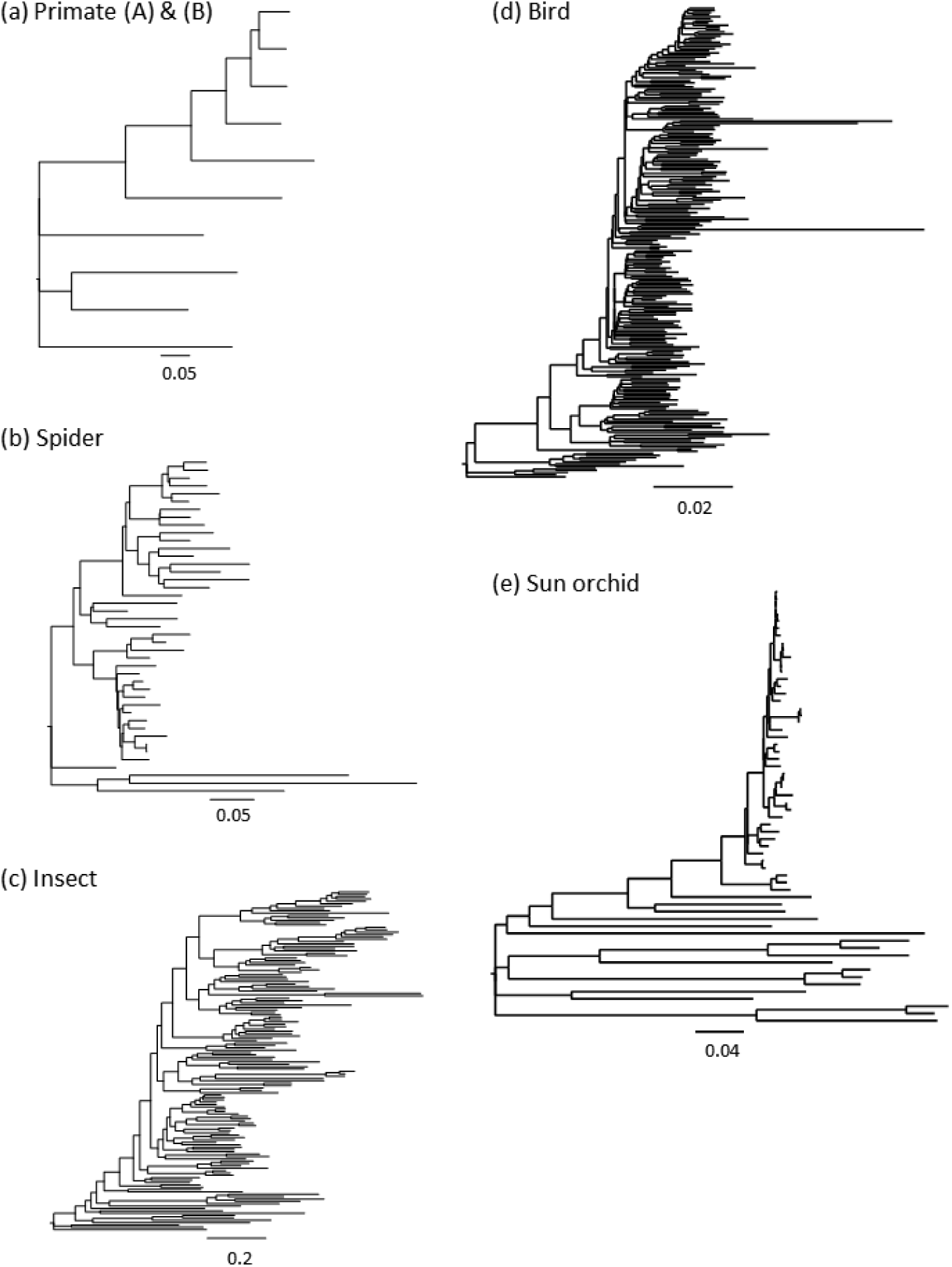
The maximum likelihood branch lengths phylogenies of empirical datasets. The dataset name refers to table 1.

